# Multi-omics approach reveals new insights into the gut microbiome of *Galleria mellonella* (Lepidoptera:Pyralidae) exposed to polyethylene diet

**DOI:** 10.1101/2021.06.04.446152

**Authors:** Samuel Latour, Grégoire Noël, Laurent Serteyn, Abdoul Razack Sare, Sébastien Massart, Frank Delvigne, Frédéric Francis

## Abstract

The current plastic pollution throughout the world implies a crucial optimization of its (bio)degradation processes. In order to identify plastic degrading bacteria and associated enzymes, the gut microbiota of insects has raised interest. Some entomological models such as *Tenebrio molitor* (L. 1758), *Plodia interpunctella* (Hübner 1813) or *Galleria mellonella* (L. 1758) have the ability to ingest and degrade polyethylene. Then, it is promising to identify the composition and the role of the gut microbiota in this process. This study takes part in this issue by investigating *G. mellonella* as a biological model feeding with a polyethylene diet. Gut microbiome samples were processed by high throughput 16S rRNA sequencing, and Enterococcaceae and Oxalobacteraceae were found to be the major bacterial families. At low polyethylene dose, we detect no bacterial community change and no amplicon sequence variant associated with the polyethylene diet suggesting microbiome resilience. The functional analysis of insects gut content was promising for the identification of plastic degrading enzymes such as the phenylacetaldehyde dehydrogenase which participate in styrene degradation. This study allows a better characterization of the gut microbiota of *G. mellonella* and provides a basis for the further biodegradation study of polyethylene based on the microorganism valorization from insect guts.

## 1. Introduction

The current plastic wastes pollution throughout the world is a serious issue. Synthetic plastics like polyethylene (PE) are used to produce packages intended for many industrial sectors such as food, pharmaceutical, cosmetics, detergent or chemical [1]. In 2016, worldwide production of plastic reached 335 megatons [2]. Indeed, its physicochemical properties made plastic a standard material in many fields. Properties such as hardness, effectiveness and cost are suitable for industries [3]. These synthetic materials have the inherent characteristic to persist in natural ecosystems and disturb it [4]. More than forty years ago, researchers have already identified a worrying amount of plastic particles drifting in the ocean [5]. Hence, impacts of micro and macro plastics on terrestrial and marine environment have been more and more tackled by scientists [6].

In order to reduce this pollution, research has been made during the last decades to enhance the degradation pathway of plastic wastes like PE. However, the high molecular weights as well as the linear carbon structure of these polymers prevent an easy and fast degradation process. Abiotic factors like heating or UV exposure allow an effective pre-ageing for a subsequent biodegradation [7]. Indeed, weathering plastic wastes are colonized by a panel of microbial colonies, typically forming biofilms on the surface [8]. Several microorganisms have been investigated to take part in the different steps of polymers biological degradation. Starting with modification of the physical and chemical properties by bio-deterioration, the size of the polymer is then reduced by enzymatic fragmentation. Then, the assimilation and mineralization allow the conversion of its molecules by the metabolism [3]. Higher organisms like insects also have the ability to participate in biodegradation processes. Among them, *Tenebrio molitor* (L. 1758), *Plodia interpunctella* (Hübner 1813) or *Galleria mellonella* (L. 1758) are strong candidates [11–13]. Recently, effects of PE on the gut microbiome of *T. molitor* larvae have been highlighted suggesting that this species can adapt its microbiome for the degradation of plastic with associated bacterial lineages [14]. The gut microbiota of insect larvae provides a new opportunity in order to improve the understanding of the plastic biodegradation pathway.

Next-generation sequencing allows a better understanding of microbial communities [15]. Sequencing of the 16S rRNA gene is a gold standard for taxonomic studies. In this study, Illumina sequencing of partial 16s rRNA sequence was applied to show the effect of plastic consumption on the midgut-hindgut microbiota of the greater wax moth caterpillars, *G. mellonella* (Lepidoptera: Pyralidae). Moreover, *G. mellonella* is most of the time considered as a model for pathogenic experimentation [16]. To date, little is known about plastic degrading enzymes especially from insect gut. Some alkane hydroxylases, isolated from bacteria, have a pivotal role to hydrolyze the PE alkanes to release primary and secondary alcohols. The digestive tract of insects could be a new opportunity to track and characterize the proteins implicated in depolymerization of waste PE [17].

Thus, examining the gut microbiome of *Galleria mellonella* larval stage is the first step for further microbiological investigations. Then, we assumed that a fraction of PE could disturb bacterial assemblages in *Galleria mellonella* gut. To test this hypothesis, the composition of the bacterial communities according to two diet approaches has been characterized: wax moth larvae only feeding on standard (STD) diet (i.e., honeybee wax) and *G. mellonella* larvae feeding on mix between PE and STD diet. Finally, we screened the enzymes expressed by bacteria and discussed their pertinence and implications into the PE biodegradation. Functional pathway of molecular determinants occurring in the microbiome and plastic degrading enzymes have been investigated based on proteomics.

## 2. Methods

### 2.1. Insect rearing and polyethylene diet

Rearing was initiated with a natural population of *G. mellonella* from Gembloux (Belgium). Temperature and humidity were maintained at 30 ± 2°C and 70 ± 10% relative humidity (RH) in darkness [18]. Honeybee wax was used to feed and host the greater wax moth population [19]. Diet methodology has been selected and optimized thanks to preliminary tests. In order to avoid premature cocoon formation, the following method was used.

We used meshed (1.5*1.5 mm) glass containers for the experiment. In each experimental structure (n=8), 5 parental larvae weighed 79 ± 26 mg were introduced. Four containers were provided with a ratio of polyethylene/honeybee wax (0.1g: 2.8g [w/w]) and four other containers were only provided with honeybee wax (control diet: 2.8g). Before first diet experiments, an entire biological cycle was elapsed (6-7 weeks) into the containers. The first two caterpillars (F1 generation) in each container, weighed more than 80 mg, were selected for dissection (n=16 larvae). Additives free low-density polyethylene was purchased from GoodFellow (ET 311351, 300×300 mm and 0.23 mm thickness).

### 2.2. Sample collection and DNA and protein extraction

Larvae were starved for one hour before dissection. Under a laminar flow, larvae were rinsed with distilled water to discard fragments. Then they were surface-sterilized with ethanol 70% for 1 minute. The dissection was carried on by immersing the whole body in 15 ml sterile PBS (NaCl 137 mM, KCl 2.7mM, Na2HPO4 10mM and KH2PO4 2mM; pH: 7.4). Larvae were maintained on the ventral side and the incision was made at the final abdominal segment with sterile scissors and scalpels. Attention was paid to keep the gut tissue intact. Finally, gut was cut through the proventricule in order to conserve the midgut and hindgut (Figure 1). Samples were then immediately transferred to a 1.5ml sterile tube and frozen in liquid nitrogen [20]. Each sample corresponds to a unique larva in order to reflect the individual midgut-hindgut microbiota. Next, disruption was made by crushing samples with sterile mortar and pestle. RLT lysis buffer (Qiagen, Hilden, Germany) was added before homogenization step. Lysate was transferred to Qiashredder column (Qiagen, Hilden, Germany). Whole DNA and proteins were extracted by the ALLprep Kit (Qiagen, Hilden, Germany), following manufacturer instructions.

**Figure 1.**
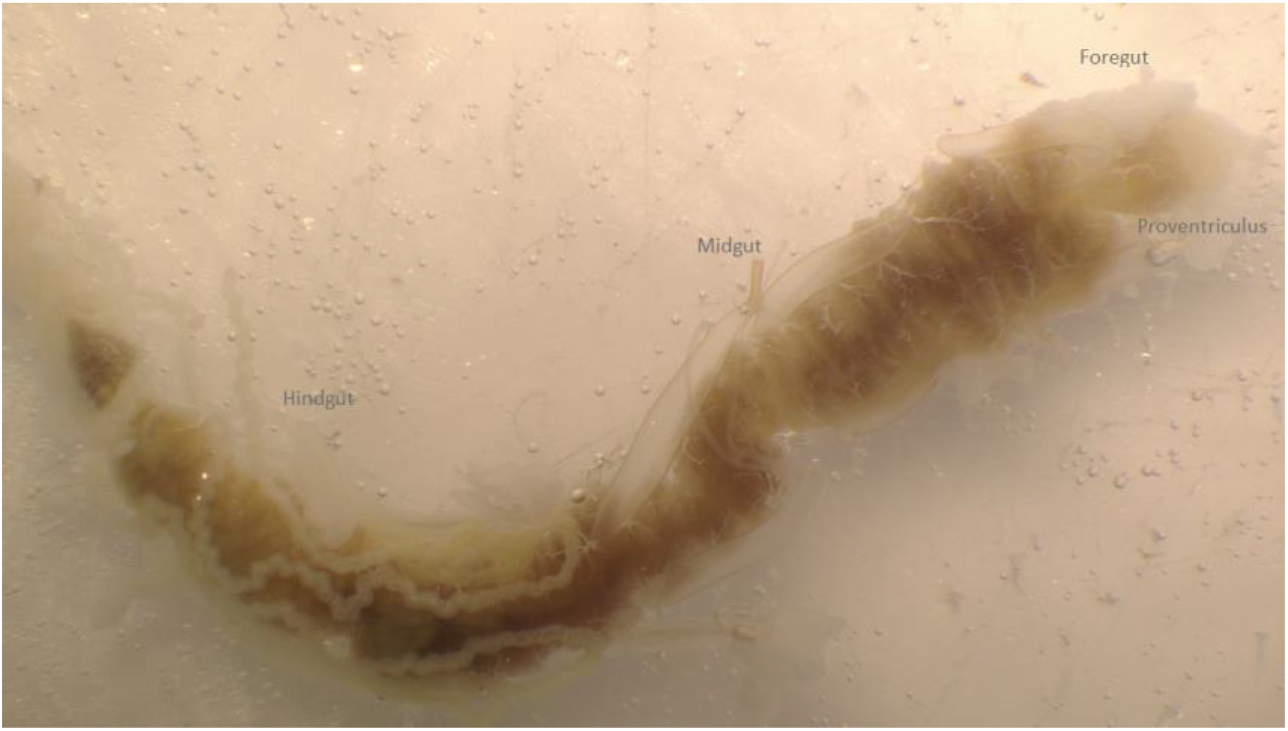
Digestive tract (mouth: top right) of *Galleria mellonella* larva dissected on the ventral side. The foregut was discarded and the midgut-hindgut part was conserved for further DNA extraction.

DNA quality from *G. mellonella* larvae was checked by Nanodrop ND-1000 spectrophotometer (NanoDrop Technologies, Wilmington, DE, USA). Double strands DNA quantity was determined by Qubit 2.0 (Quantitative dsDNA assay kit, Invitrogen, Carlsbad, California). DNA samples were kept at −20 °C until amplification.

The dry protein extract from ALLprep assay of each sample was suspended in 100 μl of rehydration buffer (7 M urea, 2 M thio-urea, 0.5% (w:vol) CHAPS). The total protein concentration was quantified by the RC-DC protein assay kit (Biorad, USA). Twenty micrograms proteins were precipited using the 2D Clean-up kit (GE, Healthcare). For each diet, four protein pellets were conserved at −80°C until further analyses. Liquid chromatography coupled with tandem mass spectrometer (LC-MS/MS) analysis by the GIGA proteomics ULiège (Liège, Belgium).

### 2.3. 16S rRNA gene amplification by PCR

The polymerase chain reaction (PCR) amplification step was performed by targeting the V3-V4 hypervariable regions of the 16S ribosomal RNA gene. Primers S-D-Bact-0341-b-S-17/S-D-Bact-0785-a-A-21, with the Illumina^TM^ (IDT, Iowa, USA) overhang adaptors in 5’ were used (5 μM) [21]. Briefly, KAPA HiFi Hot Start Ready Mix (12.5 μl) (KAPA Biosystems, Wilmington, MA, USA) was used to amplify normalized DNA samples (2.5μl, 5ng/μl). The PCR cycle program was set up as follows in a thermal cycler: 1 cycle of denaturation during 3 min. at 95 °C, followed by a denaturation cycle of 25 times 30 sec. at 95°C, an annealing step at 55°C for 30sec., then an elongation at 72°C during 30 sec. and finally, the extension was done at 72°C during 5 min. A second identical PCR was performed for samples with the control diet and four samples of the polyethylene diet to obtain enough DNA amplicons for further analysis.

### 2.4. Gene library preparation and sequencing of the 16S rRNA

Amplicon library was prepared following the 16S metagenomic sequencing library preparation protocol (Illumina, San Diego, CA, USA). Purification was performed using AMPure XP system (Beckman Coulter Genomics, Brea, CA, USA). Then, index PCR was allowed by Nextera XT index Kit (Illumina, San Diego, CA, USA). Purification was proceeding again by AMPure XP bead beating. Library was checked for quality by Agilent 2200 Tape station (Agilent Technologies, Santa Clara, CA, USA). Then, library was denatured by adding NaOH (0.2N) and the PhiX added. Finally, library was sent to DNAvision (Belgium) for Illunina MiSeq 2×250 bp paired-end sequencing.

### 2.5. High-throughput sequencing data analysis

Subsequent processing of demultiplexed raw sequences was carried out using QIIME 2 (Quantitative Insights Into Microbial Ecology) version 2018.11 software [22]. Reads were quality filtered (average min. score = 20), trimmed (F:17-245bp and R:21-245bp), de-noised, and merged thanks to DADA2 plug-in [23] resulting in high resolution of amplicon sequence variants (ASVs) for downstream assessment. The plug-in q2-sample-classifier [24] was used to the taxonomy assignment using a naïve Bayesian classifier trained on the Greengenes reference database 13.8 with 97% similarity threshold. The ensuing ASVs table was filtered from mitochondria and chloroplasts before abundance visualization and analysis with R software v3.6.3.. package ggplot2 [25] Illumina Miseq sequencing of the V3-V4 region of the 16S rRNA gene generated a total of 1,225,244 reads across 16 samples from two diets (honeycomb containing polyethylene or not), with a mean ± SD of 76,578 ± 21,867 unpaired sequences per sample. After quality filtering, 31,257 ± 9175 unique sequences were conserved for the taxonomic assignation.

In order to reduce diversity overestimate, all ASVs present in a unique sample were discarded before standardized rarefaction at 15,000 reads with GUniFrac [26]. To control for possible bias introduced by contaminants in our analyses, we applied a filtering threshold of 0.01% for low abundant ASVs as well as an additional filtering of ASVs present in less than 20% of samples. The alpha diversities were measured based on the Shannon index calculated with *vegan* [27] packages. The effect of the diet on alpha diversity was assessed by a non-parametric Wilcoxon rank-sum test. We used the Aitchison distance [28] to correct for compositional data and obtain a beta-diversity metric in an euclidean space (pseudocount of 1 added). We compared both diet (with polyethylene or not) by testing the null hypothesis that the centroids and dispersion of the groups were equal after permutational (1000 permutations) multivariate analysis of variance (PERMANOVA) with *adonis* function. Finally, to test for associations between the diet and the ASV abundances, we used a linear regression model after center-log ratio transformation (pseudocount of 1 added). We kept a false discovery rate lower than 0.05 % for all reported p-values (Benjamini–Hochberg adjustement).

### 2.6. LC-ESI-MS/MS experiments

Protein pellets were resuspended in ammonium bicarbonate 50 mM, reduced suing dithiotheotol, alkylated using iodoacetamide, and enzymatically digested using trypsin. The protein digests were purified on a Zip-Tip C18 before being injected in LC-MS/MS. The analysis was performed on an LC (Acquity UPLC Mclass – Waters) coupled with ESI-quadrupole orbitrap mass spectrometer (Q Exactive – Thermo Scientific), in positive ion mode. The obtained spectra were treated using Proteome Discoverer vs 2.1 (Thermo Scientific). Identified peptides were aligned with protein sequences of NCBI Bacteria database, using the Sequest HT server. Carbamidomethyl of Cysteines (resulting from alkylation before digestion) was set as fixed modification. Oxidation of Methionine and deamidation of Asparagine and Glutamine were set as variable modifications. Trypsin cleavage rules were applied. The proteins were filtered with the following criteria: high FDR confidence and 1 or more unique peptides.

### 2.7. Proteomic analysis

Peak intensities were log2-transformed and compared between PE and Control diets. Only protein hits that were present in three out of the four replicates of each condition were taken into account for the following quantitative proteomic analyses. T-tests between the diets were performed, using a significant threshold of 0.05 and a Benjamini-Hochberg FDR correction. Besides, proteins were considered present in a condition only when they presented valid values for at least two out of the four replicates. Finally, proteins were considered as ‘qualitatively differentially abundant proteins’ when they presented valid values in at least two out of the replicates of one condition, and in none of the replicates of the other condition. Therefore, proteins that presented a valid value in only one of the replicates of at least one of the conditions could not be taken into account in this comparative study. The proteins were annotated using the software Blast2GO (BioBam, Spain), regarding their molecular function and biological process [29].

## 3. Results

### 3.1. Taxonomic composition of the midgut-hindgut from *G. mellonella* larvae

The clustering of reads into ASV and their annotation using Greengenes database displayed Firmicutes, Proteobacteria, and Actinobacteria as the three main phyla representing respectively 51 ± 26%, 43 ± 22% and 5 ± 3% % of the 107 ASVs identified after stringent filtering from the *G. mellonella* samples (mean ± sd). The average relative abundance of the families accounting for more than 1% is shown in Figure 2A. We identified the Enterococaceae as the most abundant family whatever diet was provided. Considering the mean relative abundance for both diets, the analysis was able to identify ASVs belonging to *Enterococcus* as the main represented genus (49 ± 31 %). Then, *Curvibacter* (17 ± 10%) and the *Corynebacterium* (2 ± 1.5%) genera were among the most abundant (Figure 2B).

**Figure 2.**
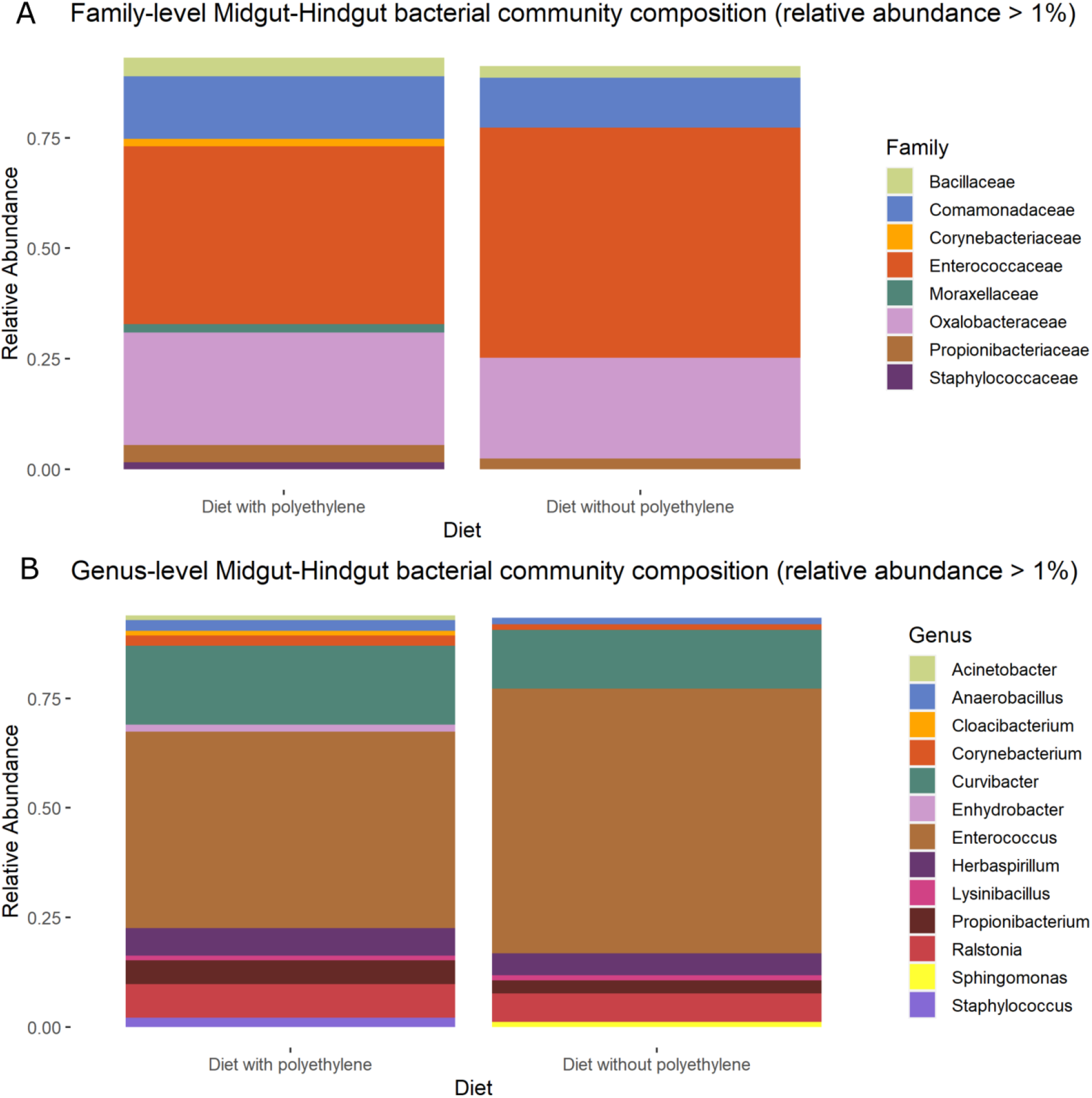
Taxonomic composition of the microbiota associated with the analyzed midgut-hindgut larval stages of *Galleria mellonella* at (A) Family level and (B) Genus level. Average abundance was calculated for honeybee wax diet (8 samples) or honeybee wax plus polyethylene diet (8 samples). Only bacteria taxa with an average relative abundance higher than 1% are reported. The others belonged to minority taxa or were not assigned.

### 3.2. Effects of polyethylene degradation on the gut microbiome

Considering the control diet (i.e. honeybee wax), Firmicutes phylum was the most abundant (56 ± 24 %). On the other hand, Proteobacteria phylum accounted for the highest abundance (47 ± 25 %) when polyethylene was present in the diet. Regarding the genus level, *Enterococcus* abundance was not significantly different between the control (57 ± 30 %) and the polyethylene diet (41 ± 33 %) and *Corynebacterium* genus was neither significantly different for the diet with polyethylene (5 ± 0.06 %) compared to the control (1 ± 1 %).

Microbial diversities have been analyzed across all two experimental diets. After removal of the low abundant ASVs (<0.01 %), features present in less than 20 % of the samples, and rarefaction to 15,000 sequences, the richness fell from 1.026 to 107 ASVs. The alpha-diversity index of the microbial community (Shannon H) was not significantly different according to both diets (Figure 3A). After controlling for compositional data structure by Aitchison transformation, we represented the variance explained by the first two eigenvectors in a principal component analysis (Figure 4). We did not see any cluster explained by the diets. This observation was confirmed by PERMANOVA (F= 1, p > 0.05) where the null hypothesis was not rejected (similarity of centroids and/or dispersion of the population based on the diet). Indeed, the distance was not significantly different for the control than the polyethylene containing diet (PERMANOVA, F=1, p > 0.05).

**Figure 3.**
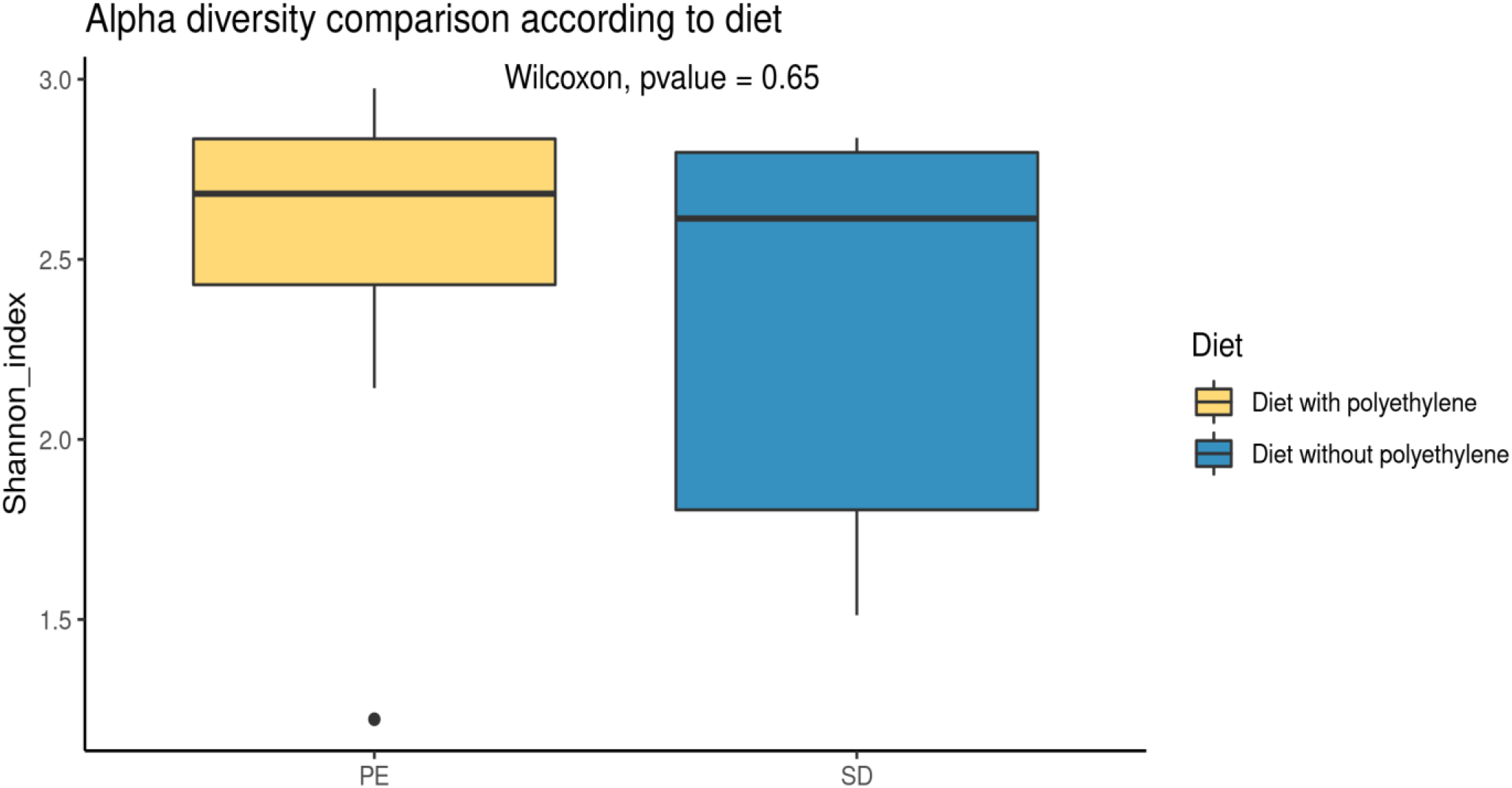
Shannon diversity index of the midgut-hindgut bacterial community from *Galleria mellonella* larval stages. Comparison is made for diet containing only honeybee wax (8 samples) or honeybee wax plus polyethylene (8 samples). Rarefaction was set at 15,000 reads depth and ASVs <0.01% or present in less than 20% of the samples were discarded. Statistical comparison of the two diets with the non-parametric test Wilcoxon-Mann-Whitney was represented for comparison of the diets.

**Figure 4.**
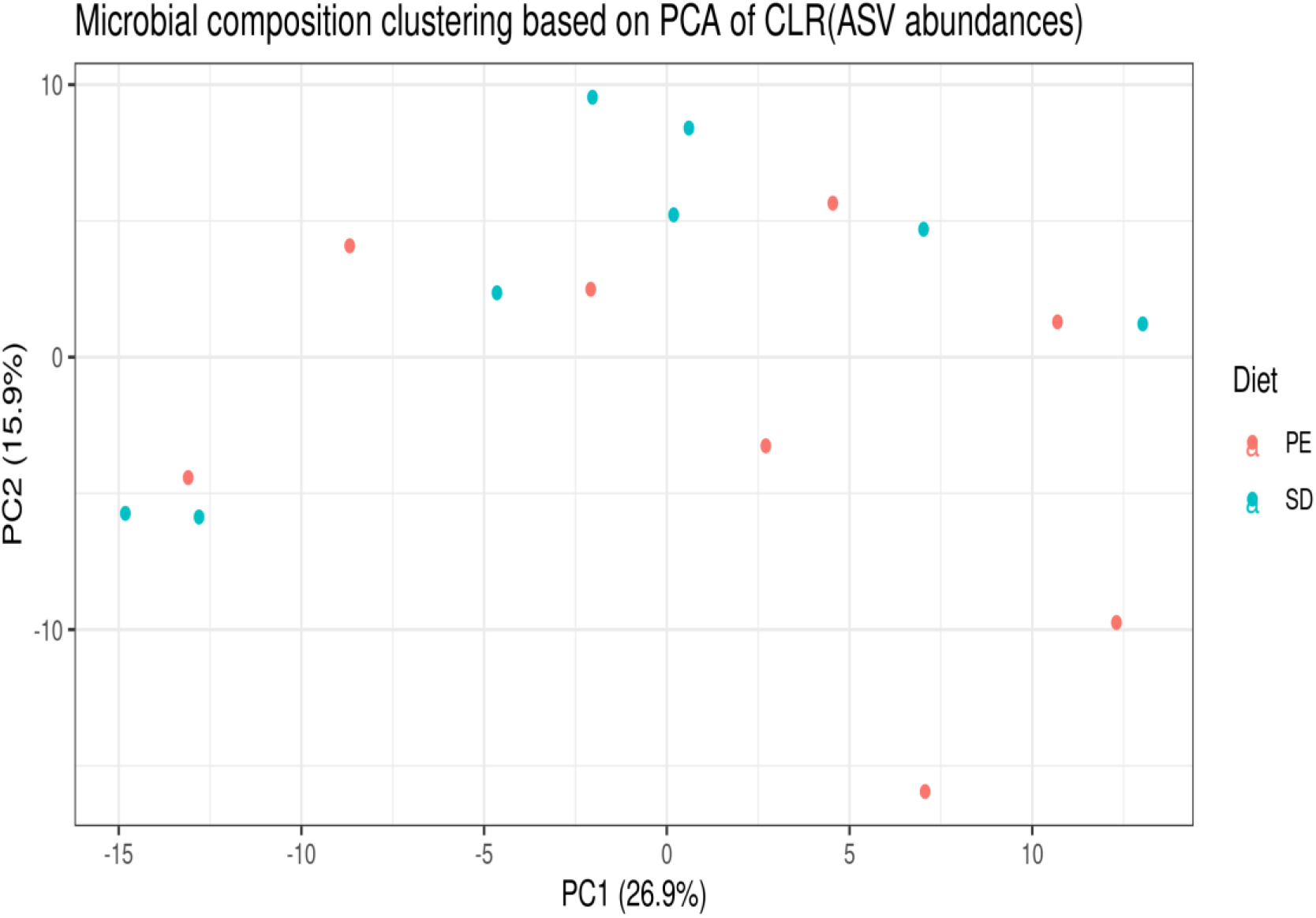
Principal Component Analysis (PCA) ordination based on the Aitchison transformed abundances (pseudocount = 1) comparing the bacterial community from *Galleria mellonella* midgut-hindgut for the diet containing only honeybee wax (SD, blue) (8 samples) or honeybee wax plus polyethylene (PE, red) (8 samples).

Linear model regression was used to assess whether specific ASVs were associated with both diets. After a pseudocount was added and abundances transformed (center-log ratio transformation), bacterial composition in the samples from larval midgut-hindgut of *G. mellonella* differed poorly between the control and the polyethylene diet. Indeed, as shown in the PCA (Figure 4), the individual variability remains high across the population and could shade the effect of polyethylene on the gut microbiota.

### 3.3. Functional insight into the larval midgut-hindgut

In order to improve the current knowledge on the protein content of the larval gut, the gel-free quantitative proteomic analysis was carried out. Among the 55 identified proteins, 34 were taken into account in a comparative analysis between the diets of *G. mellonella*, and are presented in 1. They were annotated according to their biological process, cellular component and molecular function (Figure 5). No significant discrimination were observed in the proteome of PE-fed larvae, based on the control diet, but the proteins with a fold-change higher than 1.5 were highlighted in Table 1 and presented in Figure 6. Moreover, some bacteria-associated enzymes could be attributed to putative roles in plastic degradation such as phenylacetaldehyde dehydrogenase.

**Figure 5.**
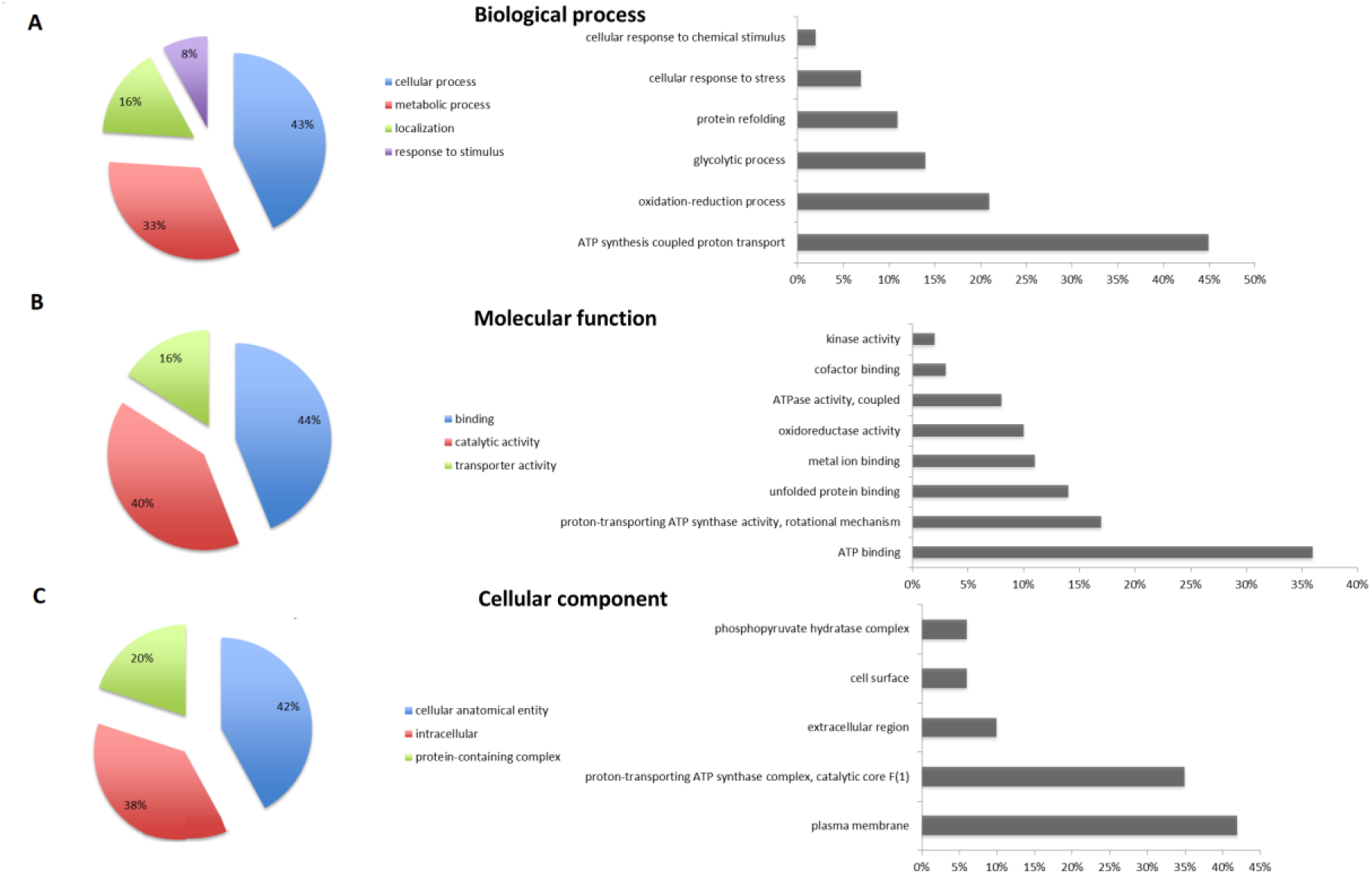
Summary of Blast2GO functional annotation of the identified proteins regarding their biological process (A), molecular function (B) and cellular component (C).

**Figure 6.**
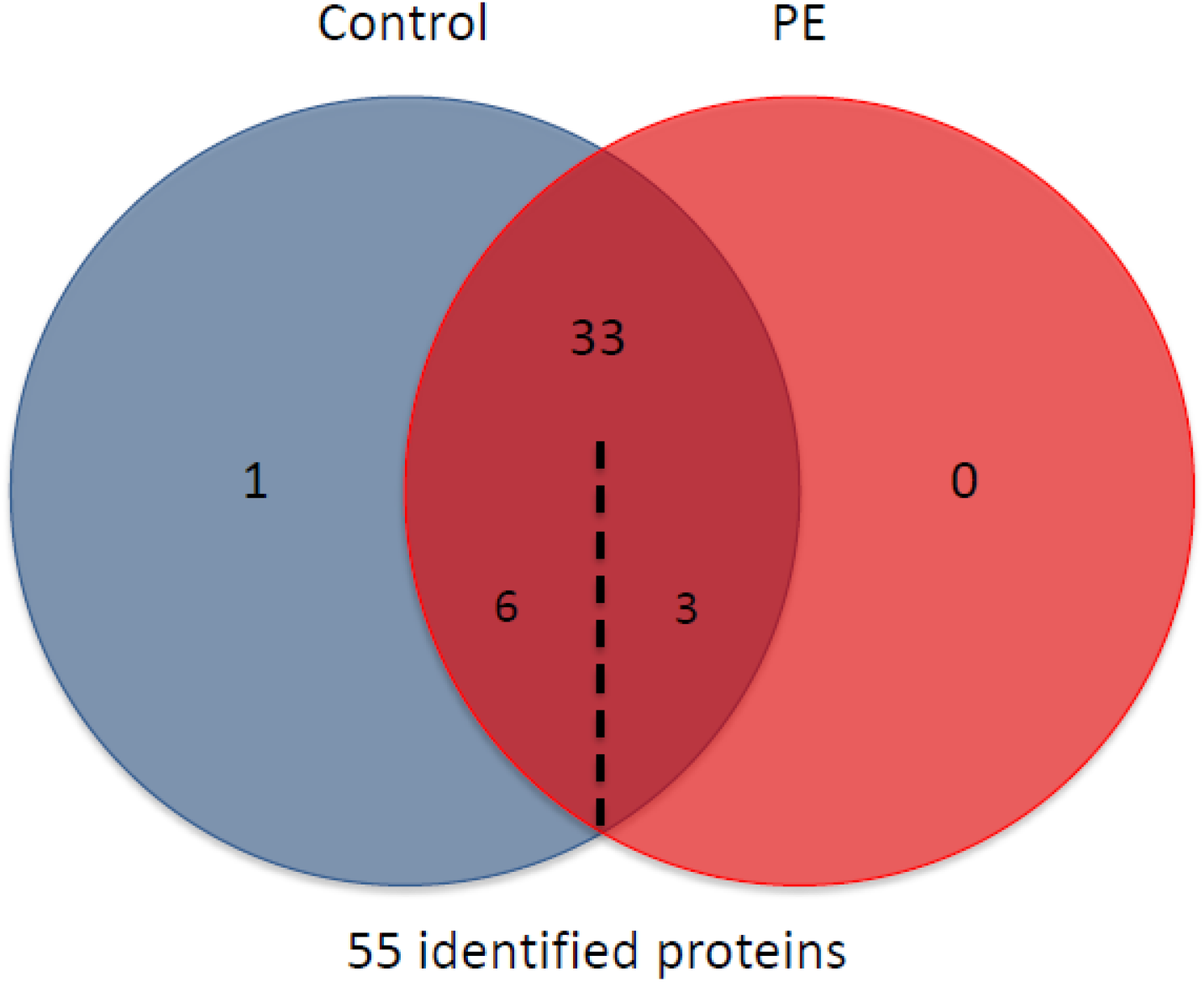
Venn diagram, distributing proteins from the larval gut of *Galleria mellonella* according to the diet treatment (“Control”: honeybee wax diet; “PE”: honeybee wax diet with polyethylene).

**Table 1.**
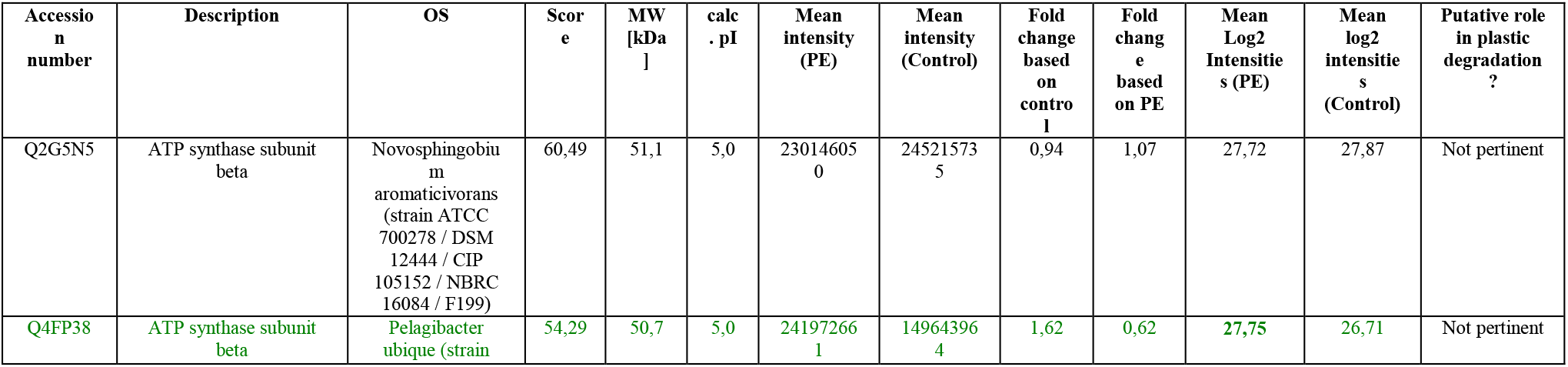
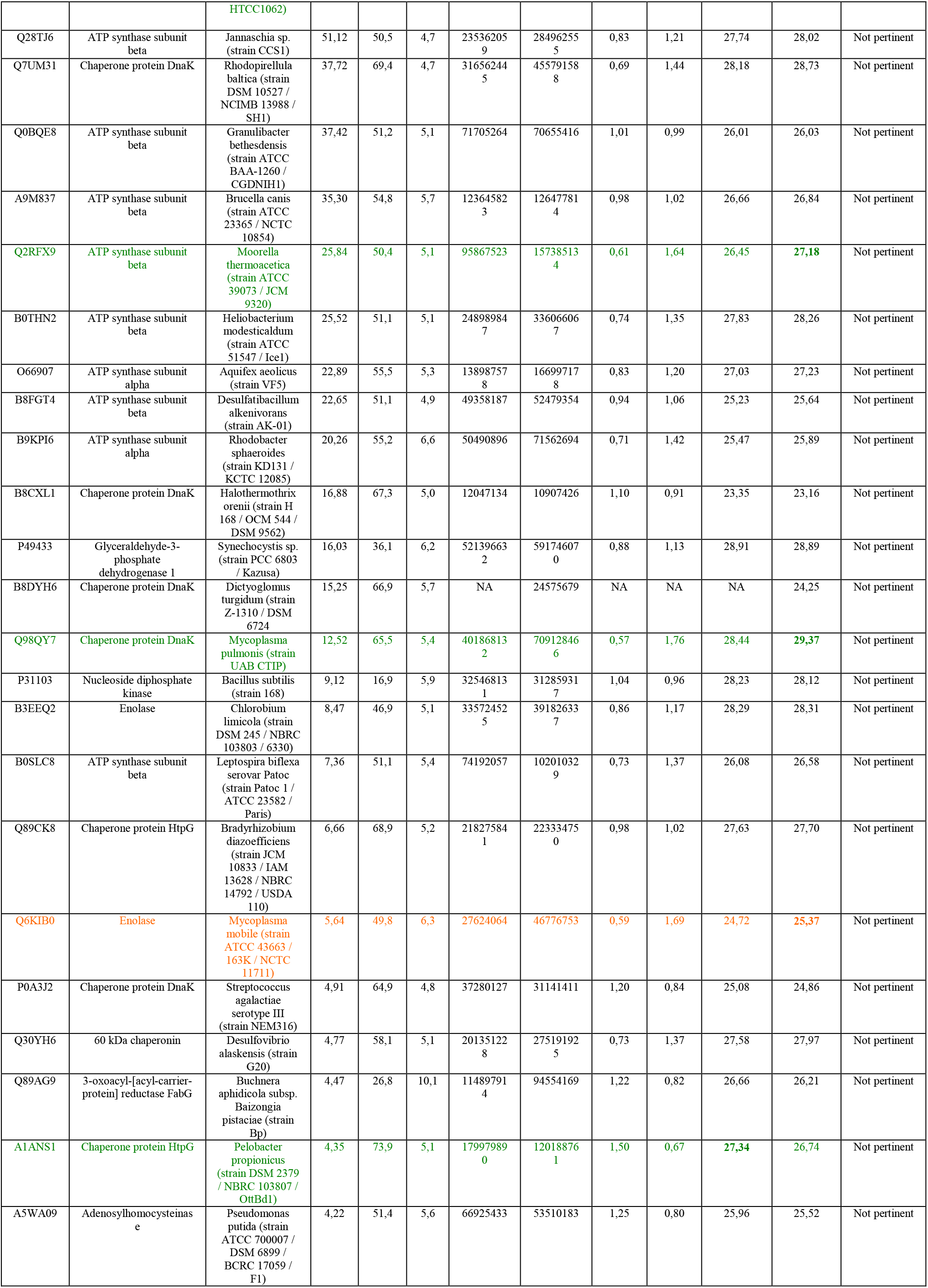
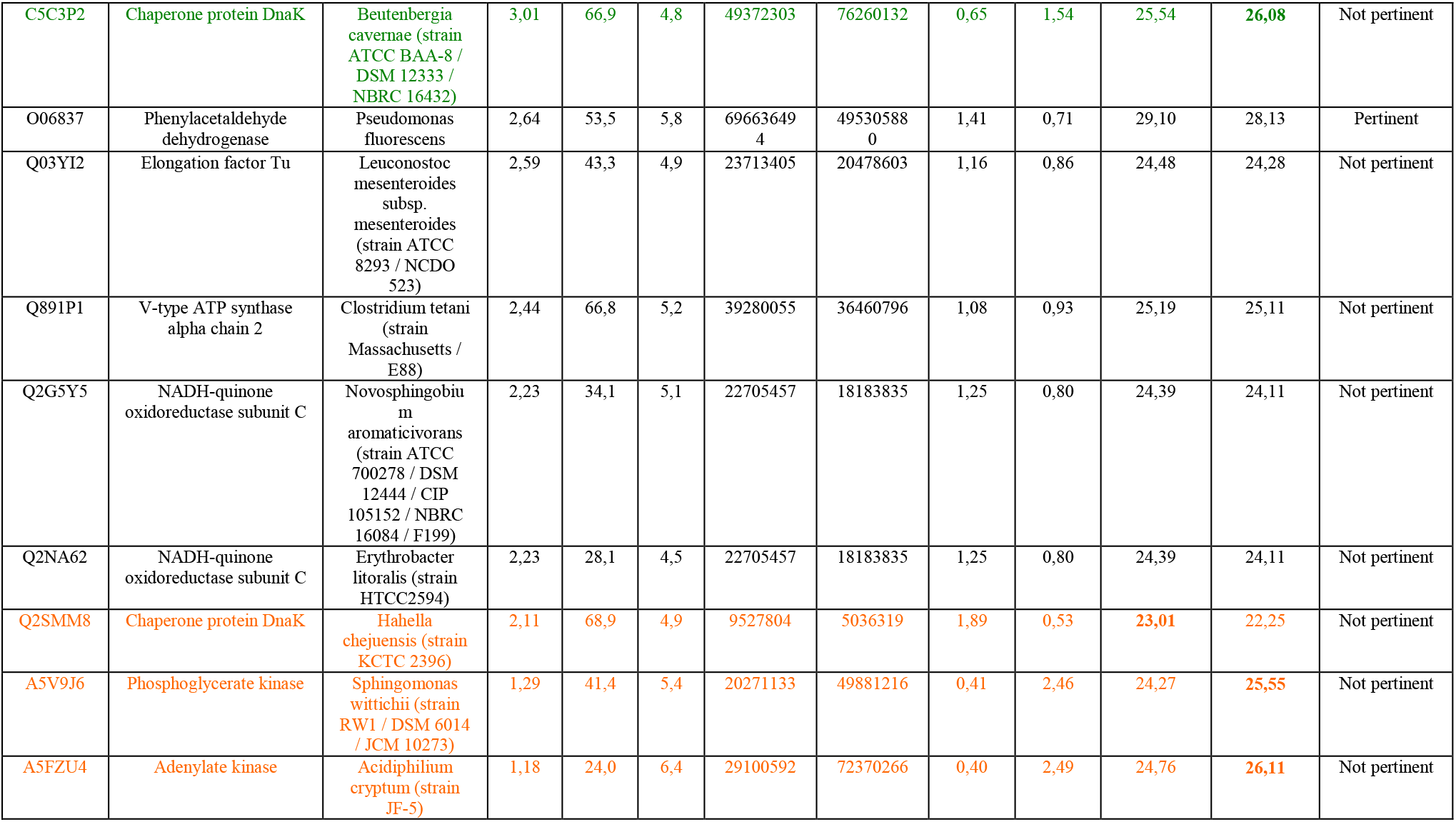
LC-MS/MS analysis of the proteins content of the midgut-hindgut from *Galleria mellonella* larvae (n=8). Only 34 proteins present in all samples were conserved. Labels highlighted in green correspond to proteins identified in minimum 2 samples of each diet treatment (or in more than 2 samples and 0 for the other) and with a fold change > 1.5. Labels highlighted in orange correspond to proteins identified in minimum 2 samples of one diet treatment but only in one sample for the other diet and with a fold change > 1.5.

## 4. Discussion

In this study, we have investigated the midgut-hindgut microbial community of the greater wax moth (*Galleria mellonella*) larvae by sequencing the v3-v4 region of the 16s rRNA gene. We also have analyzed the influence of polyethylene degradation on the taxonomic composition and phylogenetic structure of the microbiome. This study was the first to combine a genomic and proteomic approach to the midgut-hindgut content of *G. mellonella* larvae.

### 4.1. Midgut-hindgut microbiome of *Galleria mellonella* share common Insect gut pattern

The main genus present in the gut microbiome was widely shared by Lepidoptera order. Indeed, *Enterococcus* represents the highest abundance for *Spodoptera littoralis* (Boisduval 1833), *Hyles euphorbiae* (L. 1758), *Brithys crini* (F. 1775), *Bombyx mori* (L. 1758) or the moth *Plutella xylostella* (L. 1758) [30–33]. The Enterococci were highlighted to dominate the *Galleria mellonella* larvae gut due to their ability to persist despite the replacement of the epithelium, the tolerance of a high pH, the ability to degrade a wide range of carbon sources and a mutualism provided by the host [32, 34, 35]. Previous study on *G. mellonella* larvae gut states more than 80% of *Enterococcus*, however, only the midgut part of the digestive tract was used [36, 37]. Regarding the Proteobacteria, they were broadly abundant in the Pyralidae *Plodia interpunctella* [17]. It was interesting to observe that *Ralstonia*, *Curvibacter* and *Herbaspirilum* belong to three families among the most abundant in *G.mellonella* or *P. interpunctella* (Burkholderiaceae, Commamonadacea and Oxalobacteraceae). [38].

Moreover, genera *Propionibacterium* and *Staphylococcus* were among the most abundant ASVs in these two moth larvae highlighted for plastic degradation in previous studies [11, 13, 38]. The large amount of ASVs part of the Oxalobacteraceae family was not identified in any previous studies on the greater wax moth. Furthermore, these family members were commonly present in other insects gut microbiota like termites *Reticulitermes flavipes* (Kollar 1837) [39] or the sap-feeding herbivore *Mycopsylla fici* (Tryon 1895) [40]. Considering the bacterial richness, *P. interpunctella* had a higher observed OTUs. However, the diversity estimation by the Shannon index (Figure 3) was greater for *G. mellonella* suggesting a higher dominance-rarity effect of the bacterial assemblages [38]. The number of ASVs was indeed smaller but more diversified. It could be explained by the use of both midgut and hindgut for this study.

### 4.2. Effect of the polyethylene degradation on the bacterial community

Similarly to another study [37], the addition of PE to the larval diet did not have the expected effect on the bacterial communities: the alpha diversity estimates were not significantly different and PCA show no discontinuities for the two diets (Figure 3 & 4). Also, the differential abundance analysis resulted in any ASVs associated with PE. Indeed, this can be explained by the low dose of PE used in the diet which would not provide a strong enough effect to favor certain taxa. This suggests that the microbiome of *G. mellonella* larvae can be resilient to bacterial community changes at low dose of PE. The chemical composition of PE being close to honeybee wax [13], the bacteria that can potentially use it as a carbon source are present regardless of the diet.

But, we can point out some bacterial genera involving “plastic eaters” bacteria such as *Corynebacterium* genus which is present in our gut samples. This genus is gram-positive bacilli belonging to Corynebacteriaceae and frequently present in insect gut [14]. This genus was emphasized for its polyethylene degradation ability [9]. It should be interesting to deepen the identification of the gut microbiota at species level by metagenome assembled genomes (MAGs) to infer the presence of these relevant degrading bacteria. Considering that diet can shape the bacterial gut community in insect populations [41], specific bacterial populations engaged in the digestion of PE could develop (e.g. *Corynebacterium* genus). Moreover, the *G. mellonella* gut epithelial cells are able to degrade long-chain hydrocarbons such as bees wax without the presence of its gut microbiome but need its assistance in the degradation of short-chain fatty acid [42].

#### Functional insight of the midgut-hindgut proteome

According to the proteomic analysis based on bacterial dataset reference, several proteins related to respiratory chain and energy metabolic pathway were impacted by the addition of polyethylene in the honeybee wax diet. Firstly, ATP processes were identified including several ATP synthases. They are a large multi-subunit enzyme complex aiming to convert the energy of oxidation-reduction reactions of the electron transport chain to the phosphorylation of ADP. The synthesis of ATP corresponds to the respiratory chain and represents the major cell energy source by the conversion into ADP to drive cellular functions [43]. Changes in the energy metabolic pathway when changing the diets such as including new carbon source, here PE for *G. mellonella* larvae, are not surprising. Several enzymes associated to the protein folding in relation to the ATP binding molecular function were found to vary according to the plastic polymer included in the insect diet. A 60 kDa chaperonin and chaperone proteins DnaK were identified to change. Substrate binding and ATP hydrolysis by DnaK were found to be changed and are known to be regulated by chaperones [44]. In particular, the DnaK substrates include unfolded, misfolded and aggregated proteins [45]. The enzymatic cycle of DnaK alternates between ATP-bound open state and ADP-bound closed state [46]. Secondly, chaperone protein HtpG, known to display ATPase activity in cellular response to damages was found to be deregulated in *G. mellonella* feeding on polyethylene diets.

Changes in the glycolytic process and energy metabolism were observed depending on the kind of diet including polyethylene. Indeed, ATP is generated from ADP in a number of metabolic reactions of glycolysis. A glycolytic enzyme, namely phosphoglycerate kinase (PGK) was changing with insect diet. The PGK catalyzes the seventh step in glycolysis and is the first enzyme in the pathway to produce energy, rather than consume it, by the creation of two molecules of ATP from the two molecules of 1,3-bisphospoglycerate produced from glucose in earlier steps [47].

Two enolases, well known key glycolytic enzymes in the cytoplasm of prokaryotic and eukaryotic cells, were found to change according to the polyethylene occurrence in *G. mellonella* diet. Enolases are multifunctional proteins with key glycolytic role that catalyzes the switch of 2-phosphoglycerate to phosphoenolpyruvate, in the last steps of the catabolic glycolytic pathway [48]. Finally, the styrene degradation process was identified in both diets of our insect model, *G. mellonella*. Indeed, one phenylacetaldehyde dehydrogenase (PAD) was identified and known to contribute to a side-chain oxygenation that has been reported as a specific route for the styrene degradation among microorganisms [49]. Several studies revealed the attacks of the styrene vinyl-side chain found to be a specific pathway for the styrene degradation for diverse organisms, leading to the formation of the central metabolite phenylacetic acid [50].

## 5. Conclusion

This study combines a deep identification of the taxonomic composition for the gut bacterial genome and proteome from *G. mellonella* larvae. Our experiment was able to show that the *Enterococcus* genus is widely abundant in the larval stage of the greater wax moth microbiome. According to the taxonomic composition, no bacterial community differences were observed between the diets. The functional analysis of insect gut content was promising for the identification of plastic degrading enzymes, such as the phenylacetaldehyde dehydrogenase and should be deepened. Our works could provide valuable information for future plastic degradation studies. This finding coupled with the identification of *Corynebacterium* bacteria genus related to plastic degradation could have implications for discovery of potential enzymes produced by these candidate gut bacteria to degrade PE. Further investigation should be made on different polymers as well as on bacterial cultivation procedure.

## Acknowledgment

We thank Nicolas Poncelet for experimental assistance.

## Data Availability

The sequencing data have been deposited with links to BioProject accession number PRJNA730794 in the NCBI BioProject database (https://www.ncbi.nlm.nih.gov/bioproject/). The mass spectrometry proteomics data have been deposited to the ProteomeXchange Consortium via the PRIDE partner repository with the dataset identifier PXD026446.

## Code Availability

Not applicable.

